# The novel chaperone protein SRCP1 reduces insoluble SOD1 protein *in vivo* without extending ALS mouse lifespan

**DOI:** 10.1101/2020.12.21.423691

**Authors:** Ian W. Luecke, Gloria Lin, Stephanie Santarriaga, K. Matthew Scaglione, Allison D. Ebert

## Abstract

Protein misfolding and aggregation are shared features of neurodegenerative diseases, including amyotrophic lateral sclerosis (ALS), and protein quality control disruption contributes to neuronal toxicity. Therefore, reducing protein aggregation could hold therapeutic potential. We previously identified a novel chaperone protein, serine-rich chaperone protein 1 (SRCP1), that effectively prevents protein aggregation in cell culture and zebrafish models of Huntington’s disease. Here we tested whether this benefit extends to aggregated proteins found in ALS. We used viral-mediated expression of SRCP1 in *in vitro* and *in vivo* models of ALS. We found that SRCP1 reduced insoluble SOD1 protein levels in HEK293T cells overexpressing either the A4V or G93R mutant SOD1. However, the reduction of insoluble protein was not observed in either mutant C9orf72 or SOD1 ALS iPSC-derived motor neurons infected with a lentivirus expressing SRCP1. SOD1 G93A ALS mice injected with AAV-SRCP1 showed a small but significant reduction in insoluble and soluble SOD1 in both the brain and spinal cord, but SRCP1 expression did not improve mouse survival. These data indicate that SRCP1 likely reduces insoluble protein burden in a protein and/or context-dependent manner indicating a need for additional insight into SRCP1 function and therapeutic potential.

## Introduction

Amyotrophic lateral sclerosis (ALS) is a fatal, adult onset neurodegenerative disorder caused by the loss of both upper and lower motor neurons leading to paralysis, muscle atrophy, and death usually within 3-5 years of symptom onset. Most ALS cases are sporadic with no known genetic causes, but several causative gene mutations have been identified, with the most common being mutations in C9orf72, SOD1, or TDP-43 [1]. Although different genes have been associated with ALS, there is little overlap in gene function among the various mutations. Additionally, since the vast majority of ALS cases are sporadic in nature, the disease landscape is even more complicated. Yet, despite potentially divergent disease mechanisms, the presence of protein aggregates in the brain and spinal cord is consistent across both inherited and sporadic forms of ALS [2]. Protein aggregations contain a core set of proteins, including TDP-43, optineurin (OPTN), FUS, p62, and/or SOD1 [3–5], and these aggregates can induce a variety of toxic effects on cells such as inducing production of reactive oxygen species, increasing ER stress, and altering axonal transport that can lead to neuron death [6, 7]. Protein inclusions are normally prevented from forming or actively cleared by the proteostasis network, which includes molecular chaperones, ubiquitin-proteasome, and autophagy [7]. The presence of these aggregates therefore suggests an imbalance in these cellular processes and provides a targetable system that could provide therapies for ALS.

Molecular chaperones, termed heat shock proteins (Hsps), are evolutionarily conserved proteins designed to maintain protein folding in the presence of stressful cellular conditions [8]. Hsps function in an ATP-dependent and ATP-independent fashion to help refold proteins or shuttle misfolded proteins to be degraded. Recent studies have shown that Hsps are downregulated during the process of both healthy aging and neurodegeneration [9], which has a significant impact on protein clearance. In SOD1 mouse models and in post mortem ALS spinal cord samples there is evidence of decreased expression of Hsps [4] and over expression of Hsps can increase motor neuron survival *in vitro* and *in vivo* [10, 11] and delay symptom onset and slow progression in SOD1 mutant mice [12]. We previously found that motor neurons derived from mutant C9orf72 and SOD1 expressing induced pluripotent stem cells (iPSCs) exhibit insoluble protein burden and do not robustly activate the heat shock response to exogenous cellular stressors [13, 14] suggesting that ALS iPSC-derived motor neurons may lack sufficient processes to deal with insoluble protein. As such, therapies that improve protein handling without relying on activation of the endogenous heat shock response may be a valuable therapeutic option for ALS. One model organism that may have unique aspects to its proteostatic network is the cellular slime mold *Dictyostelium discoideum*. Unlike other sequenced genomes *Dictyostelium discoideum* naturally encodes roughly 10,000 homopolymeric amino acids. Among the most common repeats are polyglutamine repeats. This is interesting because polyglutamine repeats are found in a class of nine neurodegenerative diseases in which polyglutamine tracts expansion in specific genes results in protein aggregation and neurodegeneration. Previous work has demonstrated that unlike other model organisms, long polyglutamine tracts remain soluble in *Dictyostelium discoideum* [15], establishing *Dictyostelium discoideum* as an organism naturally resistant to polyglutamine aggregation.

In previous studies we used a forward genetic screen to identify a single *Dictyostelium discoideum* specific gene that is necessary and sufficient to reduce polyglutamine aggregation in cell culture, zebrafish, and iPSC-derived neurons [16]. This gene encodes an ~9kD protein, termed serine-rich chaperone protein 1 (SRCP1), that selectively recognizes and targets aggregation-prone, polyQ-expanded protein. Although SRCP1 does not contain identifiable chaperone domains, it functions through a C-terminal pseudo-amyloid domain [16]. Interestingly, data indicate that inclusions of TDP-43, FUS, and C9orf72-generated dipeptide repeat exhibit amyloid properties [17–20] suggesting that SRCP1 may target these proteins as well. Therefore, here we tested whether gene delivery of SRCP1 *in vitro* and *in vivo* would reduce insoluble protein burden and extend ALS mouse lifespan. We found that SRCP1 reduces insoluble SOD1 protein in some *in vitro* contexts but not in others. *In vivo*, SRCP1 induces a small, but significant reduction in insoluble and soluble SOD1 protein in the brain and spinal cord of ALS mice. However, this reduction was not associated with an improvement of mouse survival. Therefore, we conclude that SRCP1 may not be of direct therapeutic potential in ALS, but that SRPC1 may require specific cellular and disease protein conditions to exert positive physiological outcomes.

## Materials and Methods

### HEK293 culture and transfection

Human embryonic kidney293 (HEK293) cells were maintained in Dulbecco’s Modified Eagle’s Medium (Life Technologies) supplemented with 10% fetal bovine serum (Atlanta biologicals) and 1% Penicillin-Streptomycin (Life technologies) at 37°C and 5.2% CO2. The plasmids used in transfections were SOD1, SOD1 G85R, and SOD1 A4V all in the pEF-BOS vector. Transfections were performed with Lipofectamine 2000 (ThermoFisher Scientific) on HEK293 cells in 6-well plates at 50-70% confluency adapted from manufacturer’s instructions. Briefly, 2.5 μg of DNA per well was mixed with 150 μl of OptiMEM (Life Technologies) media and incubated for 5 minutes at room temperature. Lipofectamine 2000 in a 2:1, μl of lipofectamine: μg DNA ratio was mixed with 50 μl of OptiMEM media and incubated for 5 minutes at room temperature. DNA and Lipofectamine were mixed and incubated for 15 minutes at room temperature. Fresh media was added to cells prior to the addition of DNA and lipofectamine. Media was changed 24 hours post-transfection and cells were harvested 48 hours post-transfection. Prior to harvesting, HEK293 cells were washed three times with ice-cold 1x PBS. Samples were then lysed with 300 μl of ice-cold NETN (with protease inhibitor) and sonicated twice for 5-8 seconds.

#### iPSC Culture

iPSCs were grown on Matrigel coated 6-well plates in Essential 8 and split approximately every 7 days with Versene. Two SOD1 lines (N139K and A4V) [21], three C9orf72 lines [22], and three control lines [23–25] were used. All cells were karyotypically normal and mycoplasma free.

#### Motor Neuron Differentiation

iPSCs were differentiated into spinal cord motor neurons following a previously reported protocol (Maury et al., 2015). Briefly, motor neurons were generated by pattering EBs for two weeks in Neural Induction Medium (NIM): DMEM/F12 (1:1) (Invitrogen 11330-032), 1% N2 supplement (Invitrogen 17502-048), 1% Non-essential amino acids (11140-050), Heparin 5μg/ml (Sigma H3149), replacing medium every 3 days. On day 14, EBs were dissociated with TrypLE and plated at 50,000 cells/well onto Matrigel coated coverslips. Dissociated cells were cultured in NIM supplemented with 0.1μM Retinoic acid (Sigma R2625), 100ng/ml human sonic hedgehog (R&D 1845-SH) for one week with medium changes every 3 days. On day 21 cells were infected with a lentivirus expressing SRCP1 or GFP (MOI 5) and collected for analysis 1 week later.

#### SOD1 mice

Presymptomatic male and female SOD1 mice (SOD1-G93A, stock number: 002726) (70-80 days of age) were bilaterally injected into the cortex with 4μl of an AAV9 (10e11 vector genomes/μl) expressing either RFP (n=8) or SRCP1 (n=10). WT uninjected mice (n=3) were used as controls. Mice were euthanized at endpoint when deemed necessary due to hindlimb paralysis and/or poor body condition. The brain (cortex) and spinal cord (cervical and lumbar regions) were dissected and processed for soluble and insoluble protein analysis. Separate WT mice (n=3) were injected bilaterally with 4μl AAV-RFP (10e11 vector genomes/μl) and perfused with saline and 4% PFA two weeks later. The brain was removed and processed for immunocytochemistry to demonstrate AAV transduction.

#### Western Blot

HEK293 transfected cells were washed three times with ice-cold 1x PBS. Samples were then lysed with 300 μl of ice-cold NETN (with protease inhibitor) and sonicated twice for 5-8 seconds. Insoluble protein was separated from soluble protein in iPSCs and mouse tissues through ultra-centrifugation [13]. 50μg of sample was diluted to 100 μl with Triton X-100 lysis buffer (NaCl, 150 mM, Triton X-100, 1.0%, Tris-Cl (50 mM, pH 8.0), 1 mM EDTA, 1:100 freshly added Protease Inhibitor Cocktail Sigma P8340), for a concentration of 0.50 μg/μl. Samples were centrifuged at 45,000 rpm for 30 minutes using a Beckman TL-100 Ultracentrifuge and a TLA-45 rotor, at 4°C. Supernatants were saved as soluble protein. Pellets of insoluble protein were washed with 100μl Triton-X lysis buffer and spun at 45,000 rpm for 5 minutes for a total of 3 washes. Pellets were resuspended in 120 μl of Triton-X lysis buffer and 2% SDS. Soluble and insoluble protein were quantified with Pierce™ BCA Protein Assay Kit (23225) to ensure adequate amounts of protein. 10μg of soluble protein and 30-35μl of insoluble protein were run on 12% PAGE-SDS gel at 105 V for 90 minutes and transferred to PVDF membrane with Transfer Buffer (25 mM Trizma base, 192 mM Glycine, 500 mL of distilled water, and 20% (v/v) Methanol, pH to 8.3-8.4) at 105 V for 1 hr. For chemiluminescent detection, membranes were washed for 10 min at room temperature three times with TBST, blocked in 5% milk, and then incubated in primary antibody solution at 4°C overnight. Blots were then washed three times with TBST and then incubated in secondary antibody solution for 1hr at room temperature. Blots were then washed in chemiluminescence buffer (50mM Na_2_HPO_4_; 50 mM Na_2_CO_3_; 10 mM NaBO_3_·4H_2_O; 250 mM luminol; 90 mM coumaric acid) for film detection. For fluorescent detection, blots were washed in Odyssey blocking buffer in TBS for 1hr and primary antibody solution at 4°C overnight followed by three TBST washes and then secondary antibody incubation for 1hr at room temperature. Blots were imaged with the Odyssey Scanner (Li-COR). Primary antibodies used were rabbit anti-SOD1 (Cell Signaling 2770S, 1:1000), rabbit anti-RFP (Abcam Ab124754, 1:1000), rabbit anti-TDP43 (Proteintech 10782-2-AP, 1:1000), rat anti-HSP70 (Cell Signaling 4873S, 1:1000), mouse anti-Islet1 (Developmental Studies Hybridoma Bank, 39.4D5, 1:1000), goat anti-GFAP (Abcam Ab53554, 1:1000), goat anti-ChAT (Millipore AB114P, 1:1000), and goat anti-Iba1 (Abcam Ab5076, 1:1000). Secondary antibodies used (all from Li-Cor unless indicated otherwise) were IRDye 800 Cw Donkey anti-Rabbit (925-32213, 1:5000), IrDye 680 Donkey anti-Mouse (925-68072, 1:5000), IRDye 800 Goat anti-Mouse (925-32210, 1:5000), IRDye 800 Goat anti-Rat (926-32219, 1:5000), IRDye 680 Donkey anti-Goat (925-68074, 1:5000), and Donkey anti-rabbit HRP (Promega W401B, 1:10,000). Soluble protein samples were normalized to REVERT total protein stain (Li-Cor).

#### Immunostaining

Fixed mouse brains were sectioned on a sliding microtome (Leica) at 40μm. Sections were stained for RFP using standard procedures. Briefly, sections were washed in PBS, blocked in normal goat serum (NGS) and incubated overnight at room temperature in primary antibody solution (mouse anti-RFP (ThermoFisher MA5-15257, 1:1000) in PBS/0.3% Triton-TX, and NGS). Sections were then incubated in secondary antibody (goat anti-mouse Alexa 546 (1:500; Molecular Probes) in PBS/0.3% Triton-TX, and NGS). Sections were mounted on glass microscope slides and coverslipped using aqueous mounting media (Immunotech).

#### Statistics

Three independent *in vitro* experiments were performed for each of the iPSC lines. Western blot data for *in vitro* and *in vivo* data were analyzed using pixel intensity measurements (Li-Cor) and plotted relative to the untreated (*in vitro*) control sample or RFP vector control sample (*in vivo*). Data were analyzed using one-way ANOVA. Kaplan-Meier survival curves were analyzed using the log-rank (Mantel-Cox) test. Significance was considered with p-values < 0.05.

## Results and Discussion

In previous work we identified a novel chaperone protein in *Dictyostelium discoideum*, termed SRCP1, that had robust effects on eliminating mutant huntingtin protein aggregation in cell culture and zebrafish models of Huntington’s disease [16]. We previously found that mutant SOD1 and C9orf72 expressing iPSC-derived motor neurons exhibit increased insoluble burden of proteins known to aggregate in ALS including SOD1, optineurin (OPTN), and TDP-43 [13, 14]. We therefore asked if expression of SRCP1 could reduce protein aggregation in ALS. In initial preliminary studies (n=2) we transfected HEK293 cells with plasmids expressing wild type SOD1 or mutant G93A SOD1 or A4V SOD1 and found that co-expression of SRCP1 reduced insoluble protein accumulation for both mutant SOD1 isoforms (Fig. 1). With these encouraging data, we moved to iPSC-derived motor neurons as a more patient-specific and endogenous expression model system. We transduced control, mutant A4V and N139K SOD1, and mutant C9orf72 iPSC-derived motor neurons with lentivirus expressing either SRCP1-RFP or GFP as the vector control (n=3). Lentiviral expression of GFP in the control infected samples and RFP in the SRCP1-RFP condition were readily evident in the infected cultures (Fig. 2A,B). Consistent with our previous studies [13, 14], we found increased insoluble SOD1 and TDP-43 in the ALS iPSC-derived motor neuron cultures compared to control motor neurons (Fig. 2C). However, SRCP1 expression did not alter insoluble or soluble protein expression levels compared to GFP infected or uninfected conditions across any of the iPSC lines (Fig. 2C-E).

**Figure 1.**
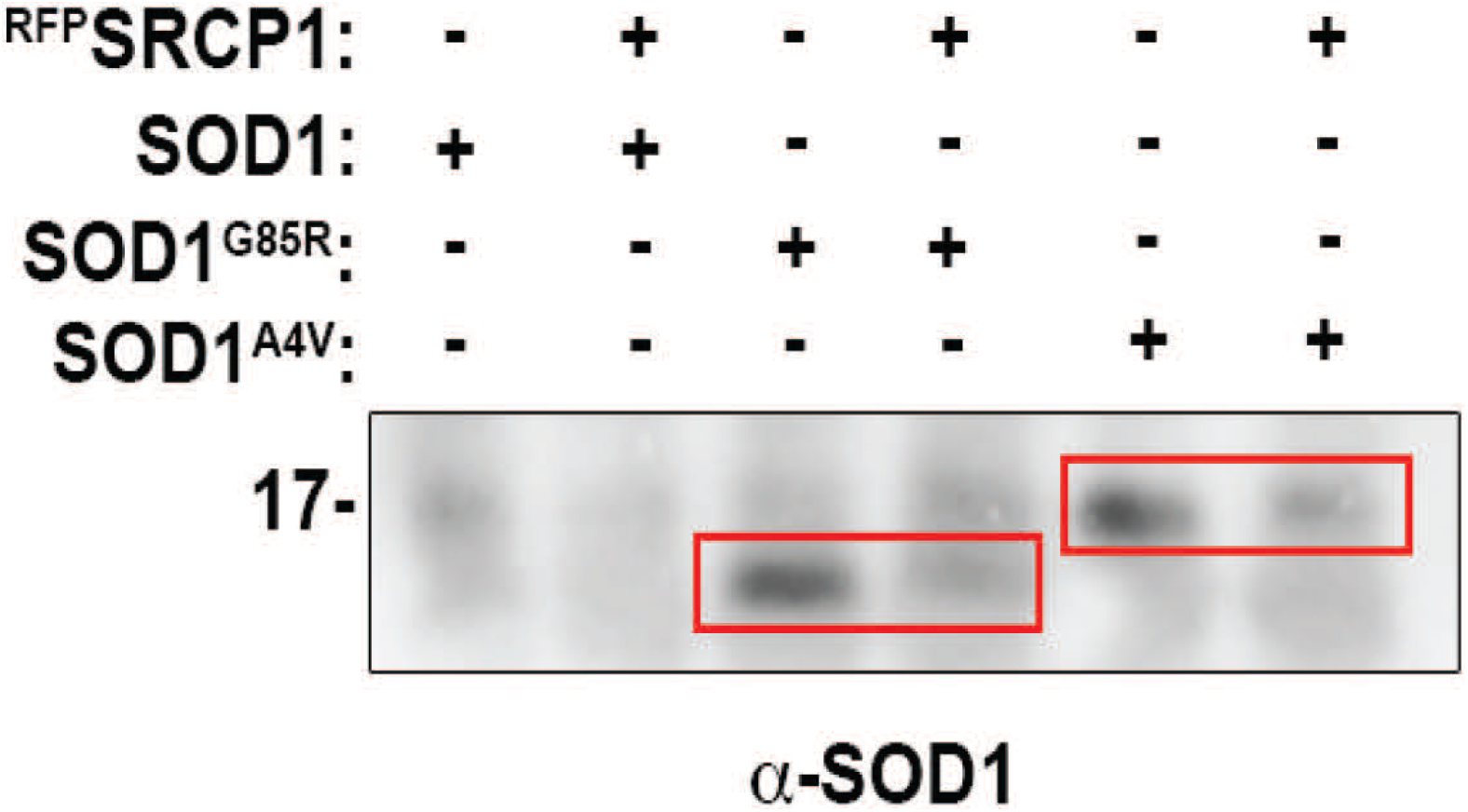
SRCP1 reduces insoluble mutant SOD1 protein. HEK293 cells expressing G85R SOD1 or A4V SOD1 showed reduced insoluble SOD1 protein upon co-transfection of SRCP1. Expression of wildtype SOD1 does not induce protein aggregation, and expression of SRCP1 had no effect. N=2

**Figure 2.**
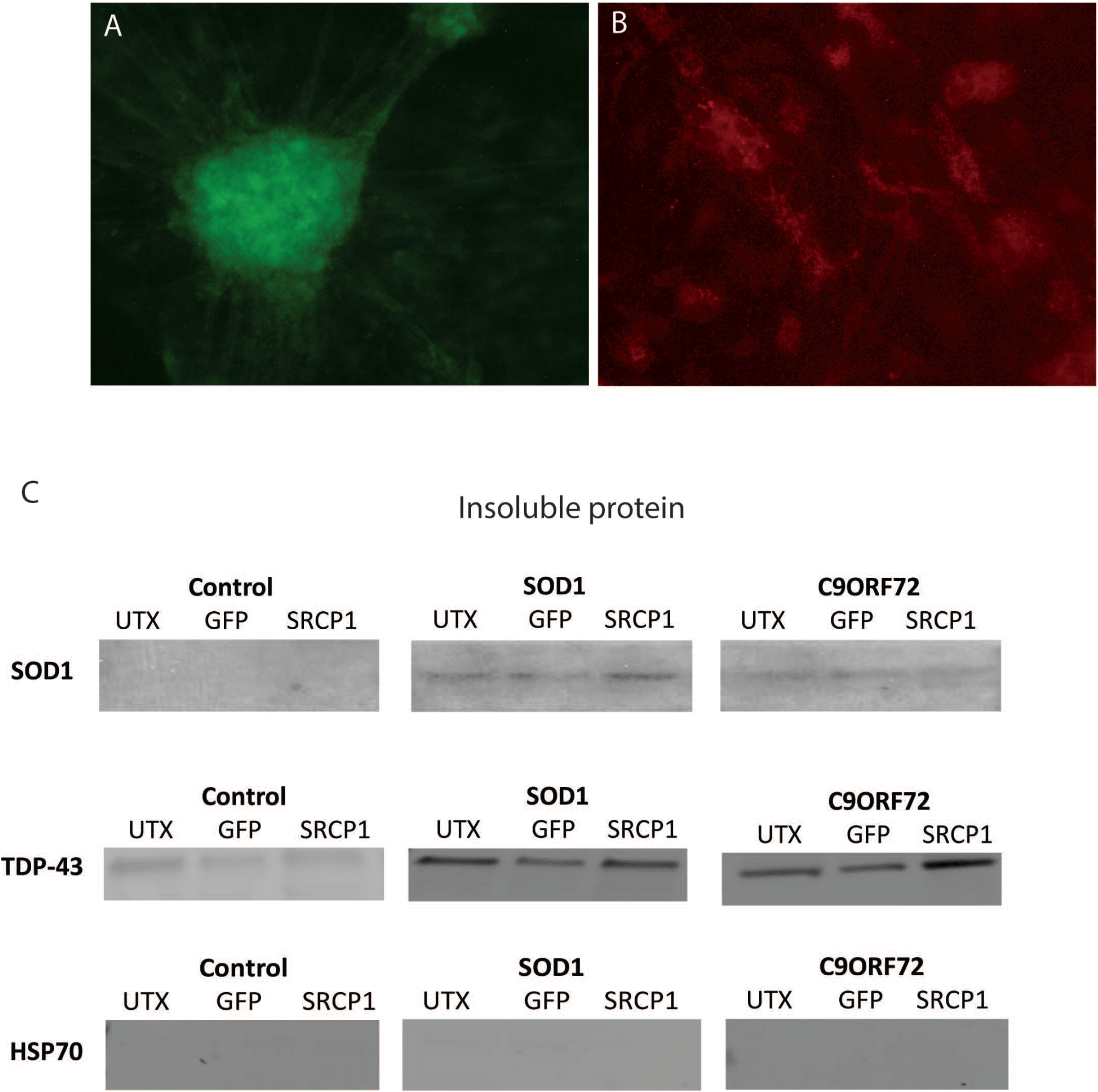
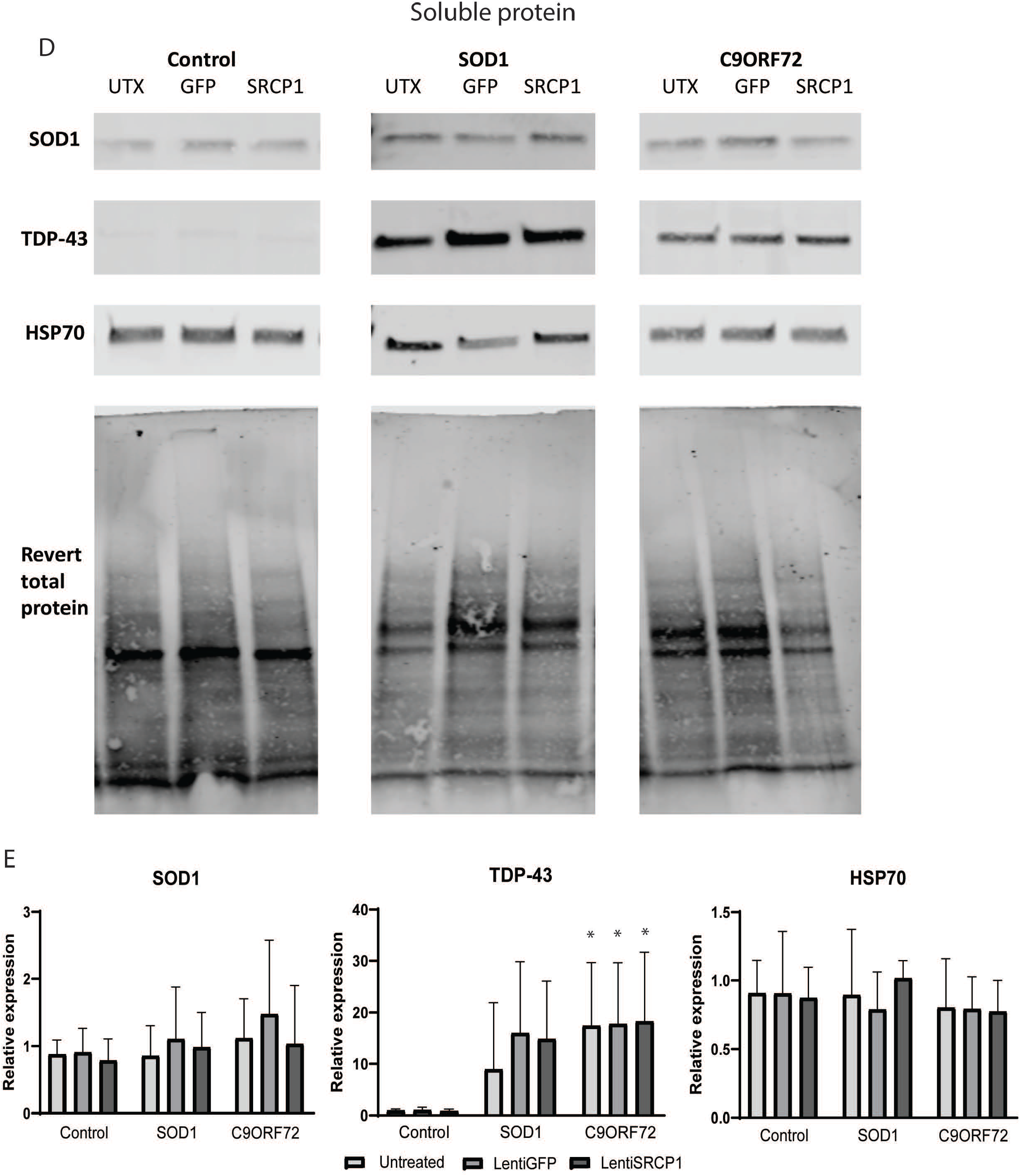
Lentiviral expression of SRCP1 had limited impact on insoluble protein burden in ALS iPSC-derived motor neurons. Live fluorescence imaging of iPSC-derived motor neurons showed transgene expression upon infection with lenti-GFP (A) and lenti-SRCP1-RFP (B). Both SOD1 and C9orf72 ALS iPSC-derived motor neurons exhibited increased insoluble SOD1 and TDP43 protein compared to control motor neurons, and lentiviral expression of SRCP1 did not reduce insoluble SOD1 or TDP-43 protein burden compared to lenti-GFP or uninfected (UTX) conditions. Lentiviral infection in healthy (control) iPSC-derived motor neurons did not alter insoluble protein burden. The lack of HSP70 expression demonstrates the efficiency of the insoluble protein purification process (C). Lentiviral expression of SRCP1 did not alter soluble SOD1 or TDP-43 protein expression in iPSC-derived motor neurons compared to lenti-GFP or uninfected (UTX) conditions. HSP70 protein expression across the samples indicates consistent soluble protein extraction. Data were normalized to total protein (D). Quantification of soluble protein showed clear increases in soluble TDP-43 protein burden in SOD1 and C9orf72 iPSC-derived motor neurons compared to control motor neurons. Neither soluble SOD1 nor HSP70 protein expression levels were different between ALS iPSC-derived motor neurons and control motor neurons (E). n=5, p<0.01 by ANOVA.

We next tested whether SRCP1 could still reduce insoluble protein burden and/or extend mouse lifespan *in vivo* despite showing mixed results in the *in vitro* models. We first showed that AAV-RFP induced robust expression 2 weeks post-injection in a WT control mouse (Fig. 3A). Next, we injected AAV-SRCP1-RFP or AAV-RFP into the cortex of presymptomatic (day 70-80) G93A SOD1 ALS mice (n=8-10) and allowed the mice to survive to endpoint. We detected RFP protein by western blot in both AAV-RFP and AAV-SRCP1-RFP injected mice at endpoint in the cortex of the brain as well as the cervical and lumbar regions of the spinal cord indicating transgene spread (Fig. 3B). Although we did not see a survival benefit from SRCP1 (Fig. 3C), we did find that there was a modest but significant reduction in insoluble SOD1 protein in the cortex of AAV-SRCP1-RFP injected mice compared to AAV-RFP injected mice (Fig. 3D,E). There was also a clear reduction in soluble SOD1 protein in both the cortex and cervical spinal cord, although this benefit did not extend to the lumbar spinal cord (Fig. 3F,G). A reduction in soluble SOD1 could be meaningful as there is evidence that soluble SOD1 may be the more toxic species [26]. Motor neuron number was not assessed here as the focus was strictly on whether SRCP1 could reduce insoluble protein. However, western blot analysis showed no consistent change in Islet1, ChAT, GFAP, or Iba1 expression in the AAV-SRCP1-RFP infected mice compared to AAV-RFP injected mice suggesting a limited impact on motor neuron survival or glial activation (data not shown).

**Figure 3.**
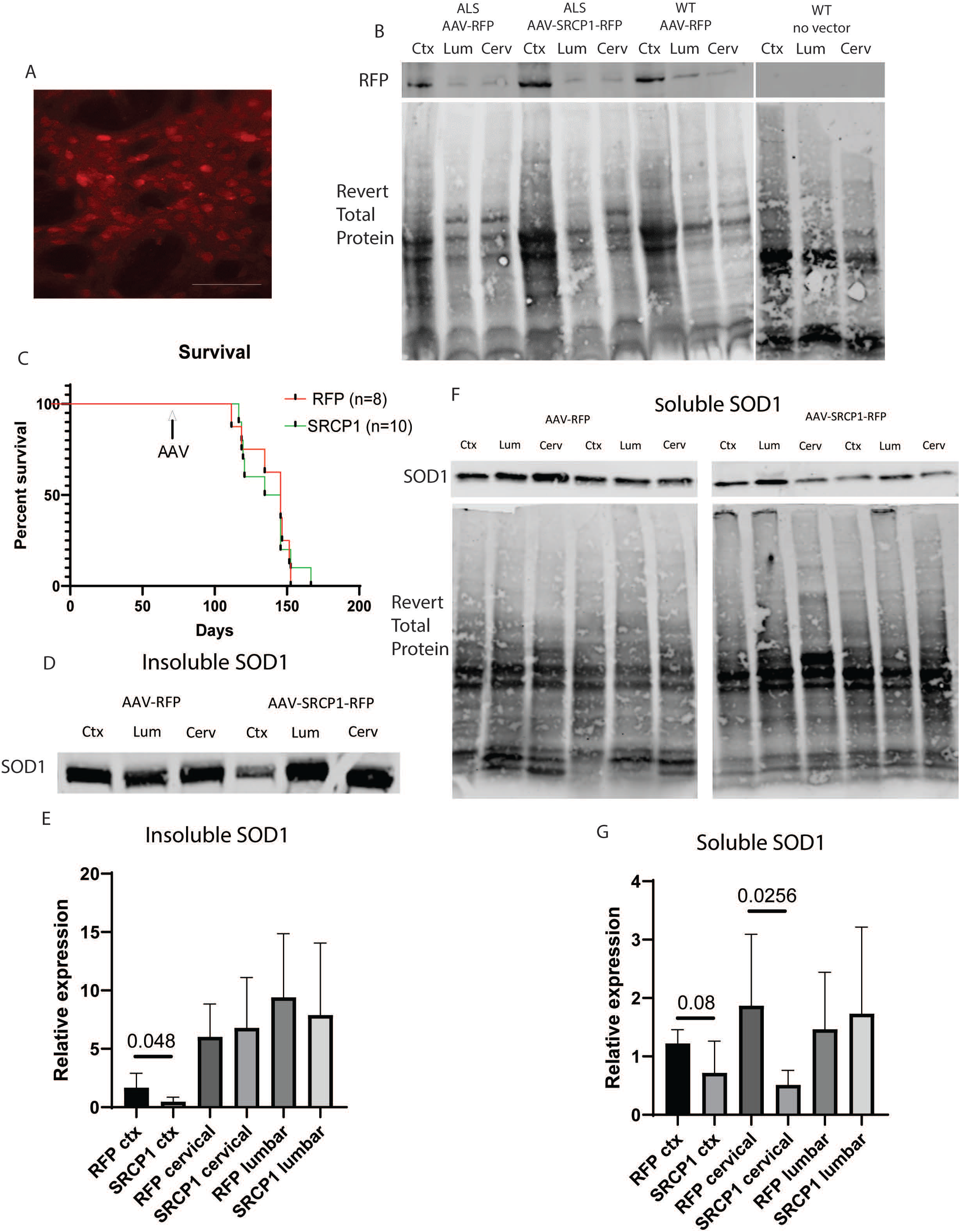
AAVSRCP1 delivery reduced SOD1 protein burden in mutant SOD1 ALS mice. Immunohistochemistry shows robust RFP expression in the brain 2 weeks post-AAVRFP injection (A). Western blot analysis shows RFP expression in the cortex as well as some expression in the lumbar (Lum) and cervical (Cerv) spinal cord indicative of transgene spread in ALS mice injected with AAV-RFP or AAV-SRCP1-RFP and a WT mouse injected with AAV-RFP. No RFP band was observed in an uninjected WT mouse (B). AAV-SRCP1-RFP did not extend ALS mouse lifespan compared to mice receiving AAVRFP (C). AAV-SRCP1-RFP significantly reduced insoluble SOD1 protein in the cortex (Ctx) of ALS mice compared to AAV-RFP injected mice but not in either the lumbar (Lum) or cervical (Cerv) spinal cord (D,E). AAV-SRCP1-RFP significantly reduced soluble SOD1 protein in the cervical (Cerv) spinal cord and showed a trend toward reduction in the cortex (Ctx) compared to AAV-RFP injected mice. AAV-SRCP1-RFP had no impact on soluble SOD1 protein in the lumbar (Lum) spinal cord (F,G).

In our previous studies testing the effect of SRCP1 in Huntington’s disease model systems, we overexpressed mutant huntingtin protein (htt) in zebrafish and control iPSC-derived neurons and found SRCP1 could reduce htt aggregations [16]. Although at the time we were unable to detect endogenous aggregated htt protein in iPSCs derived from Huntington’s disease patients, we did observe that SRCP1 expression improves neurite outgrowth [16] suggesting therapeutic benefit in the htt mutant protein expression context. In the current study we found a significant benefit of SRCP1 expression on reducing insoluble protein in the HEK cells and SOD1 mice over-expression contexts. These data suggest that SRCP1 can modulate solubility and/or degradation of mutant SOD1. However, the lack of effect on mouse lifespan could indicate that there was insufficient expression of SRCP1 to induce therapeutic benefit or that insoluble and soluble mutant SOD1 protein burden does not directly cause neuronal toxicity in the SOD1 mouse model. Additionally, because we did not observe a reduction in insoluble protein burden in conditions of endogenous mutant ALS-associated protein expression in the iPSCs, these data could indicate that protein abundance may contribute to how efficiently SRCP1 can identify and chaperone the various protein cargo. Moreover, we have previously found that ALS iPSC-derived motor neurons do not effectively modulate autophagic flux in response to increased insoluble protein burden [13], so it is possible that the benefit of SRCP1 is limited in iPSC-derived motor neurons due to disrupted proteostasis function. More research is needed to better understand the therapeutic effects of SRCP1 and its preferred protein targets, but our results suggest that viral delivery of SRCP1 may not be effective for endogenous levels of ALS-related protein cargo and will likely not be a valuable therapeutic option for ALS.

## Acknowledgements

The authors thank Dr. Emily Seminary for technical assistance with initial *in vitro* differentiations, Dr. Emily Welby for surgical assistance, and Dr. Sokol Todi for critically reading the manuscript. We also thank Dr. Christian Lorson at the University of Missouri for generating the AAV for *in vivo* experiments. The lentivirus for the *in vitro* experiments was generated by the MCW/BRI Viral Vector Core.

